# Formylglycine-generating enzyme-like proteins constitute a novel family of widespread type VI secretion system immunity proteins

**DOI:** 10.1101/2021.05.21.445229

**Authors:** Juvenal Lopez, Nguyen-Hung Le, Ki Hwan Moon, Dor Salomon, Eran Bosis, Mario F. Feldman

## Abstract

Competition is a critical aspect of bacterial life, as it enables niche establishment and facilitates the acquisition of essential nutrients. Warfare between Gram-negative bacteria is largely mediated by the type VI secretion system (T6SS), a dynamic nanoweapon that delivers toxic effector proteins from an attacking cell to adjacent bacteria in a contact-dependent manner. Effector-encoding bacteria prevent self-intoxication and kin cell killing by the expression of immunity proteins, which prevent effector toxicity by specifically binding their cognate effector and occluding its active site. In this study, we investigate Tsi3, a previously uncharacterized T6SS immunity protein present in multiple strains of the human pathogen *Acinetobacter baumannii*. We show that Tsi3 is the cognate immunity protein of the antibacterial effector of unknown function Tse3. Our bioinformatic analyses indicate that Tsi3 homologs are widespread among Gram-negative bacteria, often encoded within T6SS effector-immunity modules. Surprisingly, we found that Tsi3 homologs possess a characteristic formylglycine-generating enzyme (FGE) domain, which is present in various enzymatic proteins. Our data shows that Tsi3-mediated immunity is dependent on Tse3-Tsi3 protein-protein interactions and that Tsi3 homologs from various bacteria do not protect against Tse3-dependent bacterial killing. Thus, we conclude that Tsi3 homologs are unlikely to be functional enzymes. Collectively, our work identifies FGE domain-containing proteins as important mediators of immunity against T6SS attacks and indicates that the FGE domain can be co-opted as a scaffold in multiple proteins to carry out diverse functions.

**Importance:** Despite the wealth of knowledge on the diversity of biochemical activities carried out by T6SS effectors, comparably little is known about the various strategies bacteria employ to prevent susceptibility to T6SS-dependent bacterial killing. Our work establishes a novel family of T6SS immunity proteins with a characteristic FGE domain. This domain is present in enzymatic proteins with various catalytic activities. Our characterization of Tsi3 expands the known functions carried out by FGE-like proteins to include defense during T6SS-mediated bacterial warfare. Moreover, it highlights the evolution of FGE domain-containing proteins to carry out diverse biological functions.

## Introduction

Bacteria employ a variety of secretion systems to adapt to and thrive in the diverse environments they inhabit (1). The type VI secretion system (T6SS) of Gram-negative bacteria is an especially versatile tool implicated in various functions, including bacterial antagonism, horizontal gene transfer, metal ion acquisition, virulence, immune evasion and anti-fungal competition (2–9). This delivery device is composed of a cytosolic, membrane-anchored contractile phage tail-like complex that extends the width of the cell. When the sheath of the tail complex contracts, it propels a spiked tube structure, composed of Hcp hexameric rings topped with a VgrG trimer and a PAAR protein, from the attacking cell (predator) to an adjacent eukaryotic or prokaryotic target cell (prey). Effectors destined to secretion via the T6SS associate with the expelled Hcp-VgrG-PAAR structure via non-covalent interactions (named cargo effectors) or C-terminal translational fusions (named specialized or evolved effectors) (10).

The versatility of the T6SS is due to the diverse arsenal of effectors that is secreted by this system (11). Most T6SS effectors characterized to date mediate contact-dependent interbacterial competition, with the earliest enzymatic activities identified being peptidoglycan hydrolases, nucleases and phospholipases (12, 13). Non-enzymatic pore-forming effectors were also promptly recognized as important mediators of bacterial killing (14). More recently, the use of modern, integrative methodologies has expanded the known repertoire of antibacterial T6SS effectors to include NAD(P)+ hydrolases, ADP-ribosyl transferases, (p)ppApp synthetases, cytidine deaminases and bifunctional L,D-carboxypeptidase/L,D-transpeptidase or lytic transglycosylase/endopeptidase enzymes (15–22). Despite the wealth of information regarding T6SS effector function, comparably little is known about the diversity of strategies involved in preventing T6SS-dependent cell death (23, 24).

Mechanisms of broad protection against T6SS attacks are only beginning to be uncovered. In general, these mechanisms include modifying the T6SS effector target, preventing direct cell-to-cell contact with the bacterial predator or mounting an appropriate stress response upon perceiving an attack (25–30). Nonetheless, the most extensively studied defense mechanism against T6SS attacks is the expression of immunity proteins, which are commonly encoded in an operon with genes coding for a T6SS effector, and occasionally also a VgrG, PAAR or Hcp protein (31). Classically, immunity proteins prevent effector toxicity by specifically binding to their cognate effector and occluding its active site (32, 33). In this model, the specificity between each effector-immunity protein (E-I) pair largely limits the biological role of immunity proteins to mostly preventing kin cell killing, as immunity against non-kin effectors is dependent on the accumulation of non-kin immunity proteins (31, 34, 35).

Recent characterization of the immunity protein Tri1 from *Serratia proteamaculans* revealed a paradigm-shifting dual mechanism of immunity against T6SS attacks (15). First, similarly to all other T6SS immunity proteins reported to date, Tri1 prevents intoxication by the ADP-ribosyltransferase effector Tre1 via a mechanism of direct E-I protein-protein interactions. Additionally, however, Tri1 is a functional ADP-ribosylhydrolase capable of preventing cell toxicity by enzymatically removing Tre1-mediated ADP-ribose modifications off of the essential bacterial tubulin-like protein FtsZ (15). Importantly, this enzymatic mechanism of immunity is independent of E-I interactions, thus enabling diverse Tri1 homologs to protect against non-cognate ADP-ribosyltransferase effectors (15). Despite the clear advantage of enzymatic immunity proteins over canonical immunity proteins (i.e., broad T6SS effector protection), no additional enzymatically active T6SS immunity proteins have been described.

In this study, we investigate Tsi3, a previously uncharacterized immunity protein present in multiple strains of the human pathogen *Acinetobacter baumannii*. Our bioinformatic analyses indicate that Tsi3 homologs are widespread among Gram-negative bacteria encoding T6SSs. Surprisingly, Tsi3 homologs are predicted to structurally resemble formylglycine-generating enzymes (FGEs), which are implicated in the post-translational modification of sulfatases. Here, we employ genetic and biochemical approaches to investigate the hypothesis that Tsi3 homologs represent a novel family of enzymatic T6SS immunity proteins.

## Results

### Tse3 and Tsi3 are an E-I pair

Previously, we showed that Tse3 from Ab17978 (*ACX60_11695*, accession number WP_001070510.1) is a potent antibacterial T6SS effector of unknown function capable of killing *Escherichia coli* and *S. marcescens* (36, 37). Given the genetic proximity to *tse3*, we hypothesized that gene *ACX60_11690* (accession number WP_032046197.1, hereafter “*tsi3*”) coded for the immunity protein of Tse3 (Fig. 1a). Interestingly, *tsi3* is encoded in the opposite strand compared to *tse3*, which is an unusual genetic arrangement compared with other E-I pairs. To determine whether Tsi3 confers protection from Tse3 intoxication, we incubated wild-type (WT) Ab17978 with Ab17978*Δ3*, a mutant strain lacking the entire *vgrG3-tse3-tsi3* gene cluster (Fig. 1a), and measured Ab17978*Δ3* survival after 3.5 hours. We found that Ab17978*Δ3* was susceptible to killing by WT Ab17978 (Fig. 1b). Importantly, expression of *tsi3* in the Ab17978*Δ3* background (*tsi3+*) prevented killing by the WT strain (Fig. 1b).

**Figure 1.**
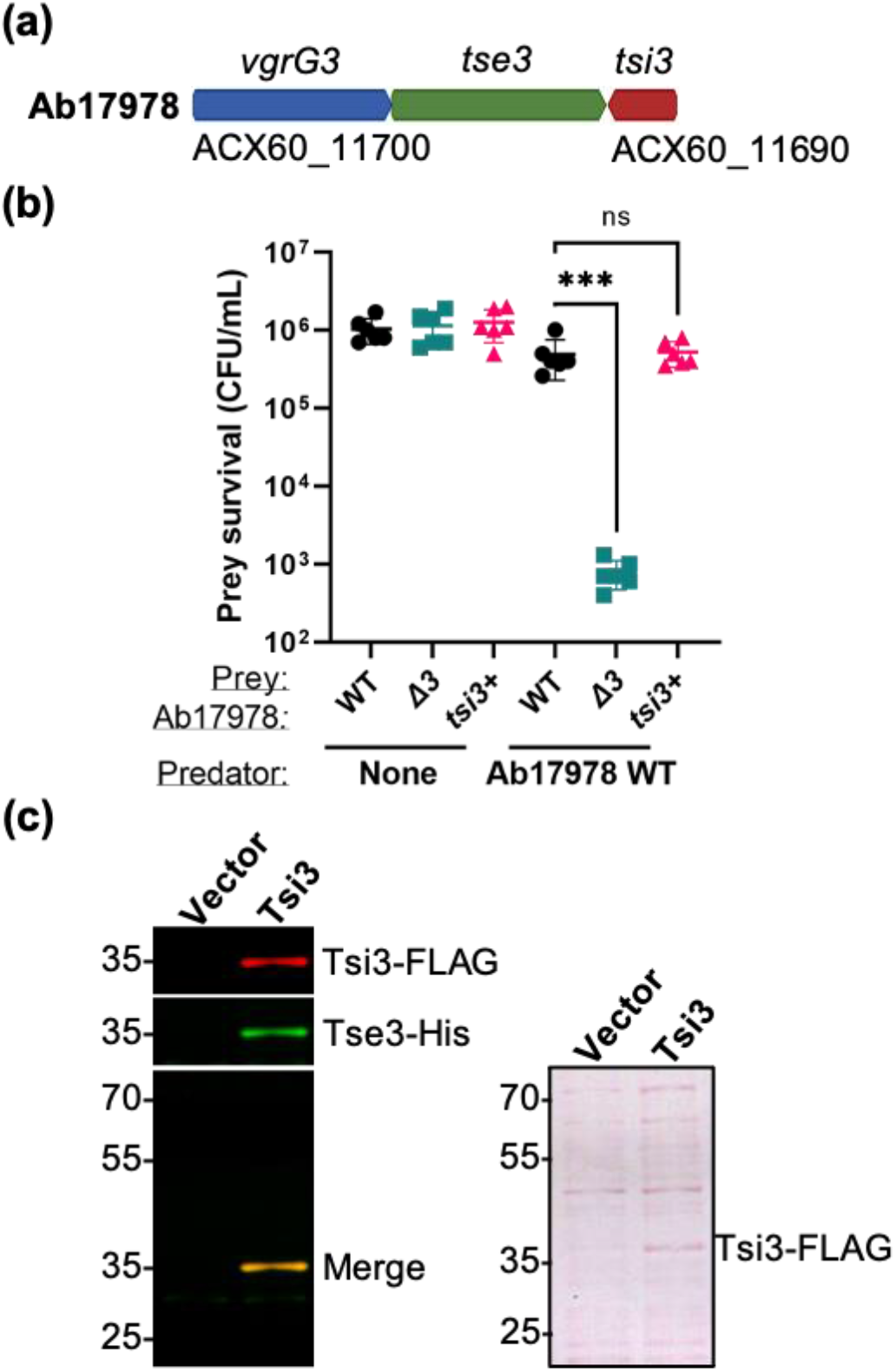
Tsi3 confers protection from Tse3 intoxication. (a) A schematic of the *vgrG3* gene cluster of Ab17978. Locus tags for the first and final genes shown are indicated. (b) Survival of the indicated prey strains following a 3.5-h incubation without a predator (None) or with WT Ab17978 predator at a 1:1 (predator:prey) ratio. Data are shown as the mean ± S.D.; *n* = 3 biological replicates in technical duplicate. ***, *P* < 0.001; ns, not significant (determined by one-way analysis of variance [ANOVA], followed by Dunnett’s multiple-comparison test). (c) Far Western blot probing for the interaction between Tse3 and Tsi3. Cell lysates from *E. coli* expressing Tsi3-FLAG or an empty vector control were separated by SDS-PAGE and transferred onto a nitrocellulose membrane in duplicate. Transferred proteins were either subjected to Far Western blot analysis (left) or to Ponceau S staining (right), which serves as a loading control.

Next, we employed a Far Western blot assay to determine whether Tsi3 physically interacts with Tse3 (Fig. S1). Briefly, cell lysates of *E. coli* overexpressing FLAG-tagged Tsi3 (Tsi3-FLAG) or a vector control were separated by SDS-PAGE and transferred onto a nitrocellulose membrane. Transferred proteins were then renatured in-membrane and incubated with purified 6xHis-tagged Tse3 (Tse3-His). After several washes, Tsi3-FLAG and Tse3-His were detected by immunoblotting. In this assay, a direct protein-protein interaction between Tsi3 and Tse3 is detected as a merge signal (FLAG and His, respectively) at the location of the nitrocellulose membrane corresponding to Tsi3. Because the nitrocellulose membrane also contains soluble proteins from *E. coli*, the absence of a Tse3-His signal elsewhere on the membrane is indicative of the specificity of the Tse3-Tsi3 interaction. Our Far Western blot assay revealed that Tse3 bound specifically to Tsi3 (Fig. 1c). Together, these results indicate that Tsi3 is the cognate immunity protein of the antibacterial effector Tse3.

### Tsi3 homologs are widespread and genetically associated with T6SS genes

We employed a bioinformatics approach (38–40) to determine the prevalence of the Tse3-Tsi3 E-I pair among diverse bacteria. Using RPS-Blast, we searched a local RefSeq database for homologs of Tsi3. We identified 3,937 Tsi3 homologs (E-value < 10^−50^) in 8,779 bacterial strains, including alpha-, beta-, gamma- and deltaproteobacteria (Fig. S2 and Dataset S1). Importantly, the vast majority of Tsi3-encoding bacteria (∼98.2%) possess a T6SS (Dataset S2). In total, we identified 26,803 occurrences of Tsi3 homologs in bacterial genomes. In ∼84.4% of cases, Tsi3 homologs were found in groups of two or more adjacently; the majority were in groups of 4 Tsi3 homologs (Fig. 2a-b and Dataset S3). Alignment of Tsi3 homologs using BLASTP revealed that in most cases, adjacently encoded Tsi3 homologs differed from each other, possibly to provide immunity against Tse3-like effectors delivered by non-kin bacterial predators (Fig. 2c and Dataset S3). Furthermore, we found that Tsi3 homologs are commonly encoded on the strand opposite to a *tse3* homolog, which is an unusual configuration for a T6SS E-I pair (Fig. 2a). Finally, genes enriched in the neighborhood of *tsi3* include those coding for T6SS structural proteins (e.g., VgrG, Hcp and PAAR), adaptor/chaperone proteins (DUF4123) and transposases, among others (Dataset S4). Our bioinformatic analyses expand on previous work by the Basler group, whose work revealed similar insights (41). Together, our results indicate that Tsi3 homologs are widespread and are often encoded within E-I modules of bacteria encoding T6SSs.

**Figure 2.**
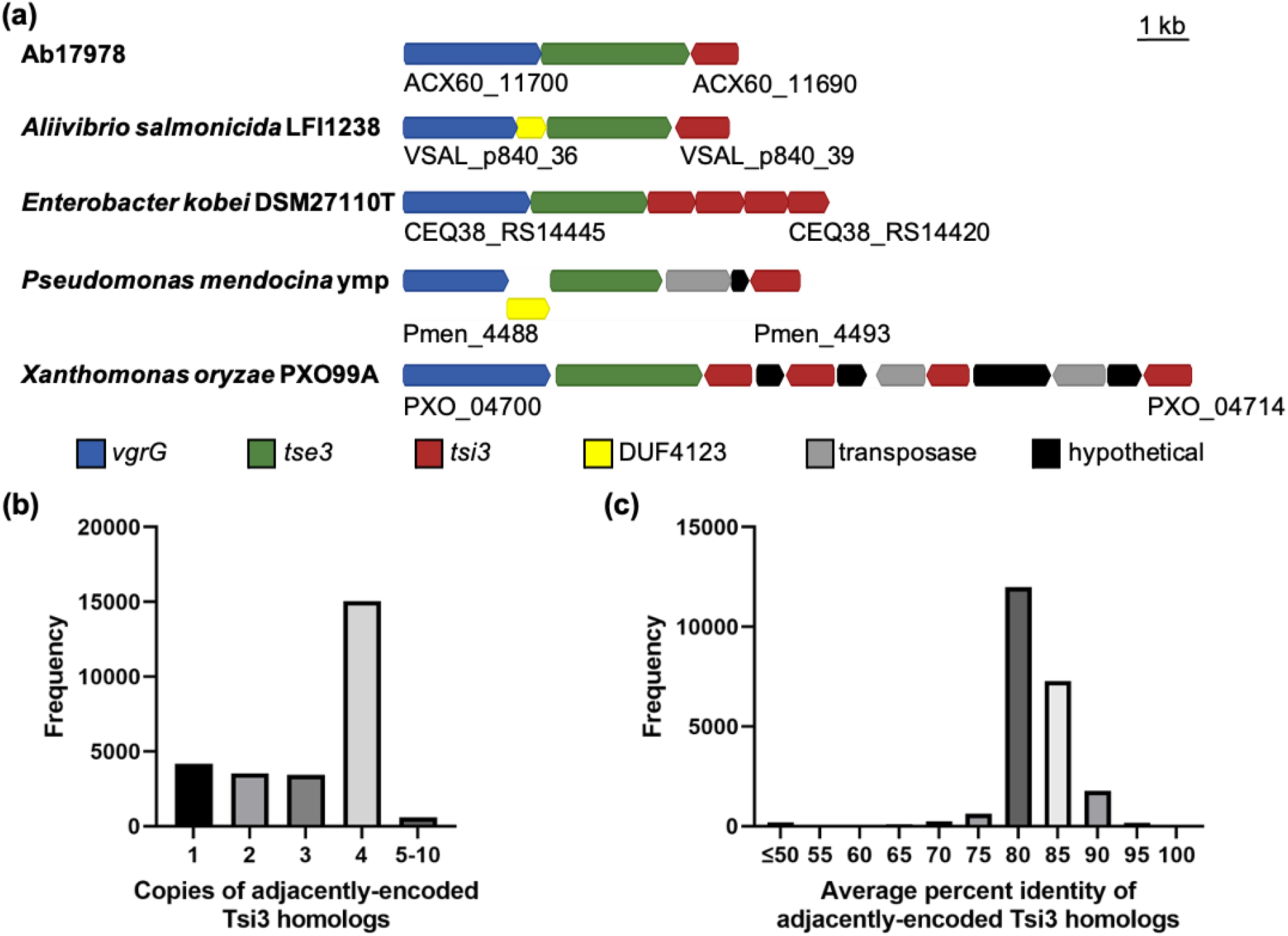
Tsi3 homologs are widespread. (a) Representative *vgrG-tse3-tsi3* loci from various bacteria. (b) A histogram showing the spread of Tsi3 homologs in groups of adjacently-encoded Tsi3 homologs. (c) A histogram showing the spread of average percentage identity among adjacently-encoded Tsi3 homologs.

### Tsi3 homologs possess a characteristic formylglycine-generating enzyme (FGE) domain

Surprisingly, domain prediction servers identified Tsi3 homologs as possessing an FGE domain (previously referred to as DUF323) (Dataset S1) (42). Besides being present in bona fide FGEs, the FGE domain is also found in non-FGE enzymes of diverse function (42), leading us to hypothesize that Tsi3 homologs function as enzymatic immunity proteins. Consistent with our hypothesis, Phyre2 identified enzymes PvdO, FGE and EgtB as the top three unique structural homologs of Tsi3 from Ab17978 (abTsi3), each with 100% confidence (Table 1). To identify putative catalytic residues of Tsi3, we modelled abTsi3 based on the known structures of PvdO from *Pseudomonas aeruginosa* (paPvdO), FGE from *Streptomyces coelicolor* (scFGE) or EgtB from *Mycobacterium thermophilum* (mtEgtB).

**Table 1.** Predicted structural homologs of Tsi3 with 100% confidence based on Phyre2.

| Rank | Protein | Fold library ID | Sequence ID |
| --- | --- | --- | --- |
| 1 | Structure of PvdO from *Pseudomonas aeruginosa* | c5hhaB_ | 21 |
| 2 | Formylglycine Generating Enzyme from *Streptomyces coelicolor* | c2q17C_ | 26 |
| 3 | Human formylglycine generating enzyme FGE | d1z70x1 | 27 |
| 4 | Formylglycine generating enzyme from *T. curvata* in complex with Cd(II) | c5nyyA_ | 26 |
| 5 | Human formylglycine generating enzyme with cysteine sulfenic acid | c1y1fX_ | 27 |
| 6 | Parologue of the human formylglycine generating enzyme | d1y4ja1 | 27 |
| 7 | Ergothioneine-biosynthetic sulfoxide synthase EgtB, apo form | c4x8bA_ | 21 |
| 8 | *Treponema denticola* variable protein 1 | c2y3cA_ | 25 |
| 9 | EgtB from *Chloracidobacterium thermophilum*, a type II sulfoxide synthase in complex with N,N,N-trimethyl-histidine | c6qkjA_ | 21 |
| 10 | Crystal Structure of CarF | c5aohA_ | 19 |
| 11 | Major Tropism Determinant P1 Variant | d1yu0a2 | 9 |
| 12 | Major Tropism Determinant I1 Variant | d1yu3a2 | 10 |
| 13 | Major Tropism Determinant P1 (Mtd-P1) Variant Complexed with *Bordetella brochiseptica* Virulence Factor Pertactin extracellular domain (Prn-E) | c2iouC_ | 14 |
| 14 | *Thermus aquaticus* variable protein (TaqVP) from diversity-generating retroelements (DGR) | c5vf4A_ | 16 |

PvdO is a putative oxidoreductase involved in the biosynthesis of the siderophore pyoverdine (43). Although the catalytic activity of PvdO has not been demonstrated *in vitro*, previous work identified a glutamate residue as essential for pyoverdine biosynthesis (43, 44). We found that abTsi3 has a glutamate residue in the equivalent position (E273) (Fig. 3a). However, this residue is not conserved among Tsi3 homologs (Fig. S3), making it unlikely to be a catalytic residue in Tsi3.

**Figure 3.**
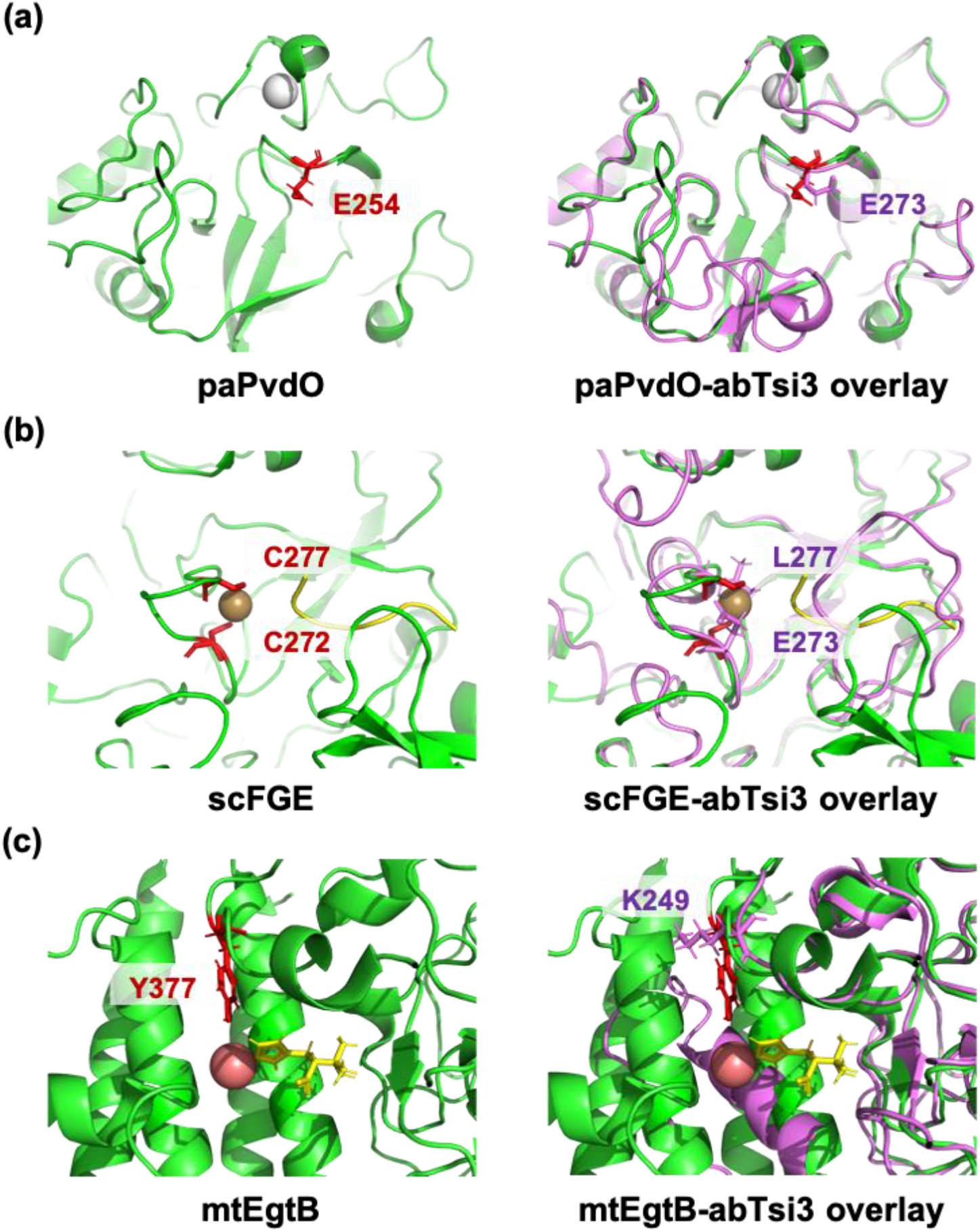
Tsi3 homologs are unlikely to be functional PvdO, FGE or EgtB enzymes. *left*, X-ray structures of (a) PvdO from *P. aeruginosa* (paPvdO, PDB: 5HHA), (b) FGE from *S. coelicolor* (scFGE, PDB: 6MUJ) or (c) EgtB from *M. thermophilum* (mtEgtB, PDB: 4×8E). Known and putative catalytic residues are shown in red and enzyme substrates are shown in yellow. The FGE substrate was extracted from PDB: 2AIK. Metal ions are shown as spheres: calcium (white), copper (brown) and manganese (pink). *right*, Overlay of known X-ray structures (green) with abTsi3 (purple) modelled to (a) paPvdO, (b) scFGE or (c) mtEgtB. Tsi3 residues replacing putative or known catalytic residues are indicated.

FGE is widespread among eukaryotic and prokaryotic organisms. FGE post-translationally activates sulfatases, which catalyze the hydrolysis of sulfate esters from various substrates, thereby carrying out diverse roles in hormone biosynthesis, nutrient acquisition, and host-pathogen interactions (45–47). Specifically, FGE converts sulfatase cysteines or serines within a conserved [C/S]-X-P-X-R motif to formylglycine (48–50). This oxygen-dependent reaction relies on two active site cysteines to coordinate a copper ion, which in turn binds the FGE substrate and primes it for reaction with oxygen (51–53). Notably, we found that Tsi3 homologs lack the catalytic cysteines essential to FGE function (Fig. 3b and Fig. S3), indicating that Tsi3 homologs are unlikely to be functional FGEs.

EgtB is a nonheme iron-dependent sulfoxide synthase involved in the biosynthesis of ergothioneine (54, 55). mtEgtB consists of an N-terminal DinB_2 domain and a C-terminal FGE domain, and the active site is composed of residues within both domains. Specifically, the FGE domain contains a metal-binding histidine triad, while the DinB_2 domain contains a catalytic tyrosine residue. The DinB_2 domain is absent in Tsi3 homologs. Thus, not surprisingly, we found that the catalytic tyrosine residue of mtEgtB is absent in abTsi3 (Fig. 3c). Moreover, no histidine triad was identified in the Tsi3 consensus sequence (Fig. S3). We conclude that Tsi3 homologs are FGE domain-containing proteins but are unlikely to be functional PvdO, FGE or EgtB enzymes.

### Non-cognate Tsi3 homologs do not provide protection against toxicity by Tse3 from Ab17978

Considering that FGE-like proteins are implicated in a wide range of activities, it remains possible that Tsi3 homologs are enzymes with an unknown catalytic activity. Based on the only report of an enzymatic mechanism of T6SS immunity (15), we premised that if Tsi3 homologs are enzymes with the same catalytic activity, non-cognate Tsi3 homologs should prevent intoxication by Tse3 from Ab17978 (abTse3). To this end, we expressed Tsi3 homologs from *A. baylyi* (ayTsi3) or *Klebsiella pneumoniae* (kpTsi3), representing 69% and 38% identity to abTsi3 (76% and 46% identity within the FGE domain), respectively, in *E. coli* and determined *E. coli* survival following co-incubation with Ab17978. Because strain Ab17978 encodes multiple antibacterial T6SS effectors, which could mask bacterial killing due to Tse3 alone, we employed strains Ab17978*ΔvgrG2,vgrG4* (hereafter Ab17978*tse3+*) and Ab17978*ΔvgrG2,vgrG4Δtse3* (hereafter Ab17978*tse3-*) as predators (36). We have previously shown that T6SS-mediated bacterial killing by strain Ab17978*tse3+* is dependent on Tse3, while the isogenic mutant strain, Ab17978*tse3-*, is unable to kill *E. coli* (36). In our assay, we also included the T6SS mutant strain, Ab17978*ΔtssM*, as a negative control for bacterial killing (Fig. 4b and Fig. S4). We found that unlike abTsi3, expression of ayTsi3 or kpTsi3 did not prevent *E. coli* killing by Ab17978*tse3+* (Fig. 4b). These results indicate that Tsi3 homologs do not provide cross-protection against non-cognate T6SS effectors.

**Figure 4.**
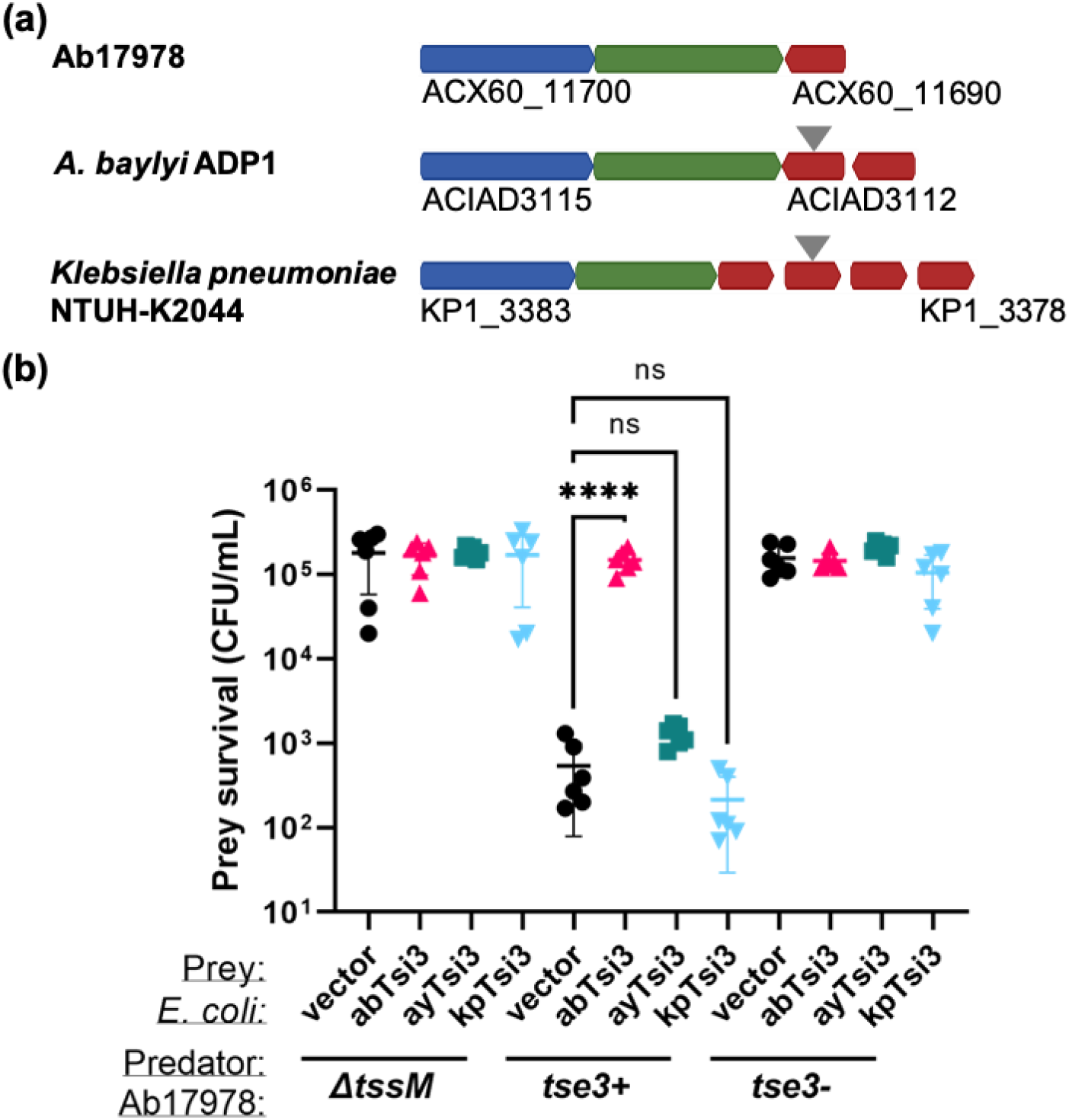
Non-cognate Tsi3 homologs do not confer immunity to abTse3. (a) Genetic context of Tsi3 homologs from various bacteria that were expressed in *E. coli* to determine functionality against abTse3. Locus tags for the first and final genes shown are indicated. The specific Tsi3 homologs expressed are denoted by gray arrows. Color scheme is consistent with Fig. 2. (b) Survival of *E. coli* prey expressing various Tsi3 homologs following a 3.5-h incubation with the indicated Ab17978 predator strains at a 1:10 predator:prey ratio. Data are shown as the mean ± S.D. from *n* = 3 biological replicates in technical duplicate. ****, *P* < 0.0001; ns, not significant (determined by one-way analysis of variance [ANOVA], followed by Dunnett’s multiple-comparison test). Ab, Ab17978; Ay, *A. baylyi* ADP1; Kp, *K. pneumoniae* NTUH-K2044.

### Disruption of Tse3-Tsi3 interaction prevents immunity to effector toxicity

Our previous results are consistent with the current paradigm that immunity proteins function by specifically binding to their cognate effector. Thus, we hypothesized that Tsi3 variants incapable of binding Tse3 will be unable to provide immunity to Tse3. To test this hypothesis, we employed our Far Western blot assay (Fig. S1) to screen several point mutants of abTsi3 for variants unable to bind abTse3 while retaining a relatively high expression level in Ab17978*Δ3* (Fig. S5). We found that mutant N194I was unable to bind Tse3, whereas mutant E236A bound Tse3 efficiently, albeit at levels lower than WT Tsi3 (Fig. 5a). In this assay, ayTsi3, which does not protect against abTse3 (Fig. 4b), serves as a negative control. Next, we tested whether the aforementioned Tsi3 variants prevent Ab17978*Δ3* killing by WT Ab17978. Unlike WT Tsi3 and E236A, expression of N194I failed to protect Ab17978*Δ3* from killing by the WT strain, suggesting that residue N194 is important for Tsi3 function. Notably, it is unlikely that residue N194 is a catalytic residue involved in a hypothetical enzymatic activity, as mutant N194A binds Tse3 and prevents Tse3-mediated toxicity (Fig. 5). Together, our results demonstrate that Tsi3-mediated immunity correlates with the ability to bind Tse3, suggesting a mechanism of immunity dependent on Tse3-Tsi3 protein-protein interactions.

**Figure 5.**
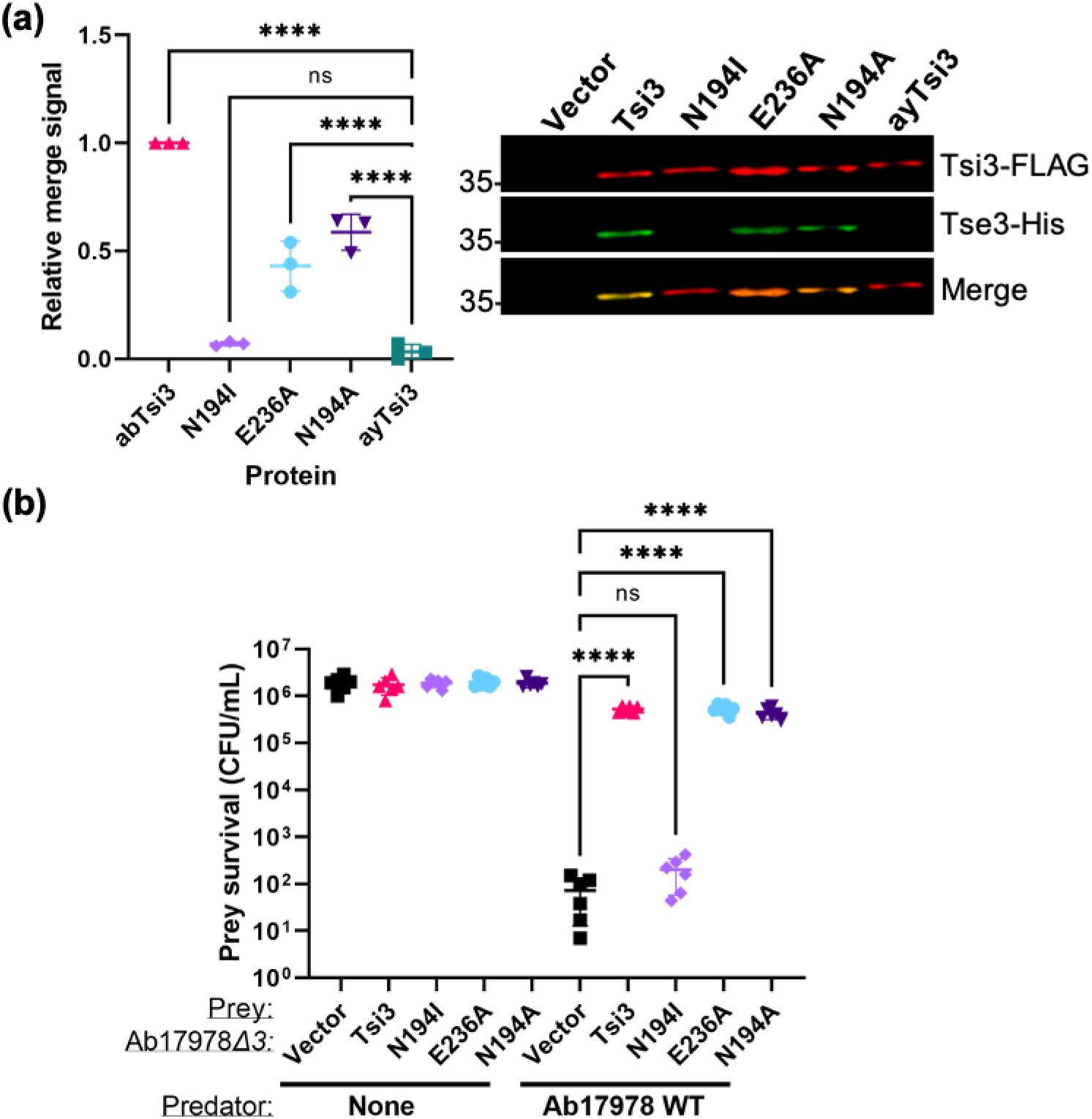
Disruption of Tse3-Tsi3 interaction prevents immunity to effector toxicity. (a) Far Western blot assay probing for the interaction between the indicated abTsi3 variants and Tse3. Data are shown as the mean ± S.D. from *n* = 3 independent experiments. ****, *P* < 0.0001; ns, not significant (determined by one-way analysis of variance [ANOVA], followed by Dunnett’s multiple-comparison test). A representative Far Western blot is shown on the right. (b) Survival of Ab17978*Δ3* prey expressing one of the indicated abTsi3 variants following a 3.5-h incubation with Ab17978 WT or no predator control at a 1:1 predator:prey ratio. Data are shown as the mean ± S.D. from *n* = 3 biological replicates in technical duplicate. ****, *P* < 0.0001; ns, not significant (determined by one-way analysis of variance [ANOVA], followed by Dunnett’s multiple-comparison test).

### Tsi3 homologs constitute a family of FGE-like T6SS immunity proteins

Our previous results indicate that although Tsi3 homologs possess an FGE domain, they are unlikely to be functional enzymes. To better understand the relationship between Tsi3 homologs and FGE domain-containing proteins, we constructed a phylogenetic tree of Tsi3 homologs and 40 representatives of the FGE family (PF03781) (Fig. 6 and Table S1). This group of proteins includes FGEs, oxygenases, serine/threonine kinases, enhancer-binding proteins (e.g., XylR), uncharacterized proteins implicated in nitrite reduction (e.g., NirV), putative oxidoreductases (e.g., PvdO), and sulfoxide synthases (e.g., EgtB) (42). Consistent with our previous results, we found that Tsi3 homologs form a clade that is distinct from that of previously characterized FGE domain-containing proteins (Fig. 6). We propose that Tsi3 homologs constitute a family of FGE domain-containing proteins specialized for mediating immunity to T6SS effectors.

**Figure 6.**
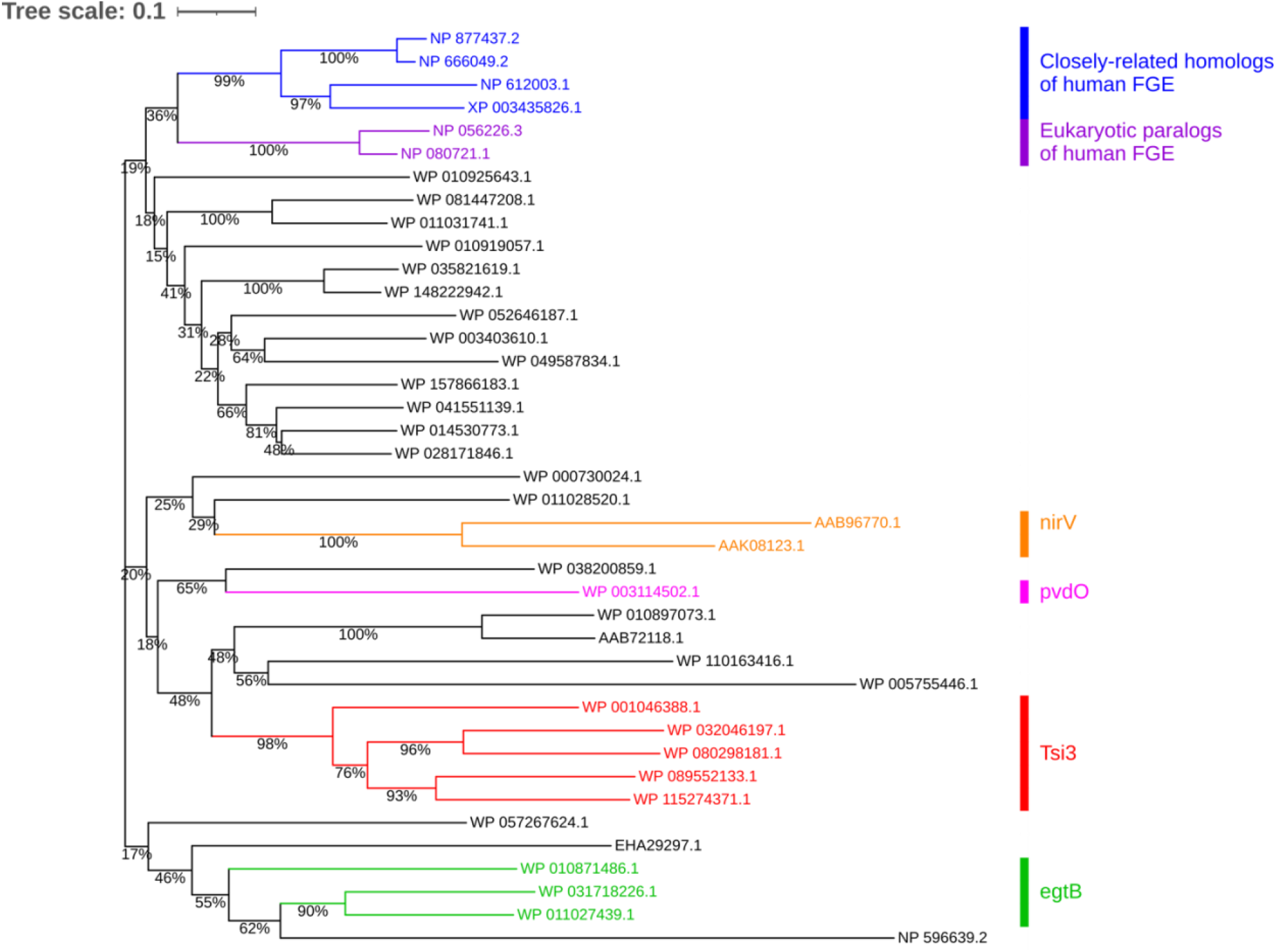
Tsi3 homologs form a clade separate from previously characterized FGE domain-containing proteins. A phylogenetic tree of representatives of the FGE family is shown. The evolutionary history was inferred using the Neighbor-Joining method. Bootstrap percentages (500 replicates) are shown next to the branches. Description of the representatives is shown in Table S1. Clades with known function are denoted on the right.

## Discussion

Since their discovery in 2010, T6SS immunity proteins were shown to prevent toxicity by interacting with their cognate effector (56). Further studies into diverse E-I pairs served to solidify this model into a paradigm for immunity protein defense (32, 57, 58). However, it was recently shown that immunity proteins can also protect potential prey via an enzymatic mechanism of immunity that is independent of E-I protein-protein interactions (15). This finding prompted us to investigate the possibility that Tsi3 homologs could represent a second family of enzymatic T6SS immunity proteins, since they possess a characteristic FGE domain, which is common in enzymes of various functions, including PvdO, FGE and EgtB. In this work, we report that Tsi3 homologs are widespread among T6SS-encoding Gram-negative bacteria, often encoded in gene clusters containing *vgrG* and *tse3* homologs. Using Ab17978 as our model organism, we experimentally determined that Tse3 and Tsi3 are in fact a cognate E-I pair and that they interact with each other. Surprisingly, although they contain a FGE domain, our structural modeling, phenotypic experiments and biochemical assays suggest that Tsi3 homologs are unlikely to be functional enzymes.

Although we cannot entirely rule out that Tsi3 homologs are enzymes, we found that Tsi3 homologs do not prevent intoxication by non-cognate Tse3-like effectors and that an abTsi3 mutant that lacks the ability to bind abTse3 is unable to provide immunity. Thus, our data suggest that the mechanism of immunity mediated by Tsi3 is dependent on the binding of the immunity protein to its cognate effector. Consistent with this proposed mechanism of Tsi3-mediated immunity, we found that in most cases, Tsi3 homologs are encoded within arrays containing more than one non-identical *tsi3* gene. It is well established that polymorphism between effector and immunity proteins underlies T6SS-dependent antagonistic bacterial interactions (59–61). Thus, the accumulation of distinct Tsi3 homologs could constitute a strategy to prevent toxicity from Tse3-like effectors of non-kin bacteria, as has been proposed for other E-I pairs in which multiple copies of immunity proteins are encoded (35, 62). For *Vibrio cholerae*, it was suggested that arrays of T6SS immunity genes are likely established through a combination of homologous and homology-facilitated illegitimate recombination (35). Further work is necessary to demonstrate the functional relevance of encoding multiple T6SS immunity genes and to provide a mechanistic understanding of the establishment of immunity gene arrays in diverse bacteria.

The striking relatedness of Tsi3 homologs to FGE domain-containing proteins suggests that these proteins likely share a common ancestor. Crystal structures from several FGE homologs indicate that the FGE domain adopts a unique “FGE fold” with low secondary structure (<20% of each β-sheets and α-helices) (49, 50, 52, 53, 63, 64). The FGE fold is also found in the X-ray crystal structures of non-FGE enzymes, including the putative oxidoreductase PvdO and the sulfoxide synthase EgtB. Our characterization of Tsi3 provides further evidence that the FGE domain can be co-opted as a scaffold in multiple proteins to carry out diverse functions. Our work expands the known functions carried out by FGE-like proteins to include defense during T6SS-dependent bacterial warfare. In addition, our findings point to the FGE domain as a potential marker to facilitate the identification of uncharacterized T6SS immunity proteins.

In sum, our work establishes Tsi3 homologs as a novel family of FGE-like immunity proteins. Future structural characterization of Tsi3 homologs in complex with their cognate effector will provide valuable information regarding the nature of the Tse3-Tsi3 interaction and may elucidate the biochemical role of the antibacterial effector Tse3.

## Materials and Methods

### Bacterial strains and growth conditions

All strains, plasmids and primers used in this study are listed in Table S2. Strains were grown in Luria-Bertani (LB) broth at 37°C with shaking. Antibiotics were added to the media when appropriate (see below).

### Generation of strain Ab17978*Δ3*

Construction of the kanamycin cassette-marked mutant strain Ab17978Δ3::Km was reported previously (65). To remove the KanR cassette, electrocompetent Ab17978Δ3::Km was transformed with plasmid pAT03 (66), which encodes the FLP recombinase. Transformants were plated on LB agar containing 2 mM IPTG supplemented with carbenicillin (200 µg/mL). Removal of the kanamycin resistant cassette was confirmed by PCR and sequencing.

### Generation of expression plasmids for Ab17978*Δ3* and *E. coli*

The pWH1266-based (67) construct for abTsi3-His expression was generated by linearizing pWH-*vgrGi*-6xHis (68) by PCR, amplifying *tsi3* from genomic DNA of Ab17978, then ligating both PCR products by In-Fusion (Takara Bio, Mountain View, CA), according to the manufacturer’s instructions. Point mutants were generated using the QuikChange II site-directed mutagenesis kit (Agilent Technologies, Santa Clara, CA), according to the manufacturer’s instructions. The specific alanine mutations made to Tsi3 were selected to represent a wide sample of charged and polar residues conserved among Tsi3 homologs. Mutations N194I and R287C were selected because mutations in equivalent residues of human FGE were shown to disrupt FGE function (49). Ab17978*Δ3* transformants were selected for using 15 µg/mL tetracycline. These strains were employed as prey in bacterial killing assays.

The pBAVMCS-based (69) constructs for abTsi3-His expression wes generated by PCR amplification of the abTsi3 gene and restriction cloning into BamHI/PstI sites. Due to homology between the two *tsi3* homologs encoded by *A. baylyi* ADP1, we first amplified a segment of DNA containing both *tsi3* homologs. Then, we digested that fragment with EcoRI to separate both homologs. The segment containing gene ACIAD3113 was then amplified, digested, and ligated into pBAVMCS at KpnI/SalI sites (ACIAD3113 has internal BamHI/PstI sites). The gene encoding the *tsi3* homolog from *K. pneumoniae* NTUH-K2044 was obtained as a geneblock from Integrated DNA Technologies (IDT), Coralville, Iowa. This geneblock served as a template for PCR amplification, restriction enzyme digestion, and ligation into the BamHI/PstI sites. The specific *A. baylyi* ADP1 and *K. pneumoniae* NTUH-K2044 Tsi3 homologs selected for this study have the highest percentage identity to abTsi3 in their respective E-I module. *E. coli* Rosetta 2 transformants were selected for using 50 µg/mL kanamycin and 12.5 µg/mL chloramphenicol. These strains were employed as prey in bacterial killing assays.

abTse3-His was cloned into vector pET28a by In-Fusion (Takara Bio). *E. coli* Rosetta 2 transformants were selected for using 30 µg/mL kanamycin and 12.5 µg/mL chloramphenicol. abTsi3-FLAG and ayTsi3-FLAG were cloned into vector pETDUET at sites NdeI/KpnI. Point mutants of abTsi3 were generated as described above. *E. coli* Rosetta 2 transformants were selected for using 200 µg/mL ampicillin and 12.5 µg/mL chloramphenicol. These strains were used to perform Far Western blot assays probing for the interaction between Tse3 and Tsi3 (see below).

All constructs were verified by PCR and sequencing.

### Bacterial killing assays

Overnight cultures of predator and prey strains were washed three times in fresh LB and normalized to an OD600 of 1. Predator and prey strains were then mixed at the appropriate ratio (1:1 WT Ab17978:Ab17978*Δ3* or 1:10 WT Ab17978:*E. coli* Rosetta 2, respectively), spotted onto dry LB-agar plates, and incubated for 3.5 h at 37°C. Spots were then resuspended in 1 mL LB and serially diluted onto dry LB-agar plates supplemented with antibiotics appropriate to select for surviving prey. Plates were incubated overnight at 37°C and CFUs were quantified thereafter.

### Tse3 purification

Overnight cultures of *E. coli* Rosetta 2 cells harboring pET28a-Tse3-6xHis were diluted 1:100 into autoinduction media ZYM-5052 (70) and grown for 16 h at 30°C with the appropriate antibiotics. Cells were harvested by centrifugation, resuspended in resuspension buffer (50 mM Tris, 300 mM NaCl, 25 mM imidazole, pH 8) containing EDTA-free protease inhibitor tablets, and lysed by three passages through a cell disruptor at 35 kpsi. Cell lysates were clarified by centrifugation and loaded onto a Ni-NTA agarose column (Gold Bio, St. Louis, MO) equilibrated with resuspension buffer. The column was then washed with resuspension buffer and wash buffer (50 mM Tris, 300 mM NaCl, 50 mM imidazole, pH 8), and immobilized Tse3 was eluted with elution buffer (50 mM Tris, 300 mM NaCl, 300 mM imidazole, pH 8). Purified Tse3 was concentrated, buffer exchanged into 50 mM Tris, 150 mM NaCl, pH 8, and concentrated once more to a final concentration of ∼ 1 mg/mL.

### Induction of FLAG-tagged abTsi3 variants and ayTsi3

Overnight cultures of *E. coli* Rosetta 2 cells expressing C-terminal FLAG-tagged Tsi3 variants/homologs were diluted to an OD600 of 0.05 in fresh LB containing the appropriate antibiotics. The cultures were then grown at 37°C (shaking) until mid-exponential phase and induced with 1 mM IPTG for 4 h at 30°C. Next, cells were harvested by centrifugation and resuspended in resuspension buffer.

### Far Western blot assay and immunoblotting

The interaction between Tse3 and Tsi3 variants was determined by Far Western blot using a previously published protocol with few modifications (71). First, resuspended *E. coli* Rosetta 2 cells expressing C-terminal FLAG-tagged Tsi3 or variants thereof (see above) were treated with concentrated lysis buffer to a final concentration of 1% triton X-100, 100 μg/mL lysozyme, 100 μg/mL DNaseI, 10 mM MgCl_2_, 10 mM CaCl_2_ and EDTA-protease inhibitor tablet. Cells were lysed by sonication and clarified lysates were loaded onto a 12% SDS-PAGE gel (in duplicate) and ran at 150 V. Proteins were then transferred at room temperature (RT) onto a nitrocellulose membrane using a semi-dry transfer cell (Biorad) (20 V, 35 min) and cold transfer buffer (10% methanol, 0.1% SDS, 25 mM Tris, 200 mM glycine). One membrane was treated with Ponceau S stain to visualize all cell lysate proteins, and the other membrane was used for Far Western blot analysis.

For the Far Western blot, transferred proteins were denatured and renatured in the nitrocellulose membrane by treatment with decreasing concentrations of guanidine-HCl then incubated overnight at 4°C without guanidine-HCl, as previously described (71); however, instead of 2% milk for each treatment, we used Odyssey Blocking Buffer (TBS) (LI-COR, Lincoln, NE) at up to 50% by volume. The membrane containing renatured proteins was then blocked with Odyssey Blocking Buffer (TBS) at room temperature for 1 h, treated with purified Tse3-His in “protein-binding buffer” (71) at 1 μg/mL and incubated overnight at 4°C. Unbound Tse3-His was removed by washing with Tris-buffered saline–Tween (TBST). Proteins were then detected with monoclonal mouse anti-FLAG M2 (1:1000; Sigma-Aldrich, St. Louis, MO) and polyclonal rabbit anti-6xHis (1:2,000; Invitrogen, Waltham, MA) as well as IRDye-conjugated anti-mouse and anti-rabbit secondary antibodies (both at 1:15,000; LI-COR Biosciences). Blots were visualized with an Odyssey CLx imaging system (LI-COR Biosciences). Quantification was done using Image Studio 5.2 (https://www.licor.com/bio/help/imagestudio5/index.html#Introduction_help.html%3FTocPath%3D_____2), as previously described (72). The data are presented as relative merge signal, where the His/FLAG value corresponding to WT Tsi3 is defined as 1.

### Structural modeling of abTsi3

Structural homologs of abTsi3 were identified using Phyre2 (73). Three independent structural models of abTsi3 were generated using I-TASSER (74) based on the known X-ray crystal structures of paPvdO (PDB: 5HHA), scFGE (PDB: 6MUJ) and mtEgtB (PDB: 4×8E). The FGE substrate was extracted from PDB: 2AIK. All structures were visualized using the PyMOL Molecular Graphics System, Version 2.3.4 (Schrödinger, LLC).

### Identification of Tsi3 homologs using PSI-Blast

Identification of Tsi3 homologs was performed as described previously for other proteins (38–40). First, the PSSM of Tsi3 was constructed using the amino acid sequence of Tsi3 from Ab17978 (WP_032046197.1). Five iterations of PSI-BLAST (75) against the reference protein database were performed. In each iteration, a maximum of 500 hits with an e-value threshold of 10^−6^ were used. RPS-BLAST (75) was then used to identify Tsi3 homologs in a local RefSeq database (downloaded April 17^th^, 2021). E-value threshold was set to 10^−50^.

The sequences of proteins located in the neighborhood of Tsi3 homologs were analyzed; conserved domains were identified (see below), signal peptides and cleavage sites were predicted using SignalP 5.0 (76), and transmembrane topology and cleavage sites were predicted using Phobius (77). Identification of conserved domains enriched in Tsi3 genomic neighborhoods was performed as described before (78).

The neighborhood of Tsi3 homologs was scanned both upstream and downstream for the existence of adjacently encoded Tsi3 homologs. Two Tsi3 homologs were counted as adjacently encoded proteins if not more than one unrelated protein was identified in between the proteins. Tsi3 homologs were aligned using BLASTP and average percent identity of the Tsi3 homolog to the adjacently encoded Tsi3 homologs was calculated.

### Illustration of conserved residues in Tsi3 homologs

Tsi3 homologs were aligned using Clustal Omega (www.ebi.ac.uk/Tools/msa/clustalo/) (79). Aligned columns missing from Ab17978 Tsi3 (WP_032046197.1) were removed from the alignment. WebLogo was created using the WebLogo 3 server (weblogo.threeplusone.com/) (80).

### Identification of conserved domains and additional domains

The Conserved Domain Database (CDD) version 3.19 and related information were downloaded from NCBI (81). RPS-BLAST was employed to identify conserved domains and the output was processed using the Post-RPS-BLAST Processing Utility v0.1. The expect value threshold was set to 10^−5^.

### Identification of T6SS core components in bacterial genomes

RPS-BLAST was employed to identify the T6SS core components in bacterial genomes containing Tsi3 homologs, as described before (39). Briefly, the proteins were aligned against 11 COGs that were shown to specifically predict T6SS (COG3516, COG3517, COG3157, COG3521, COG3522, COG3455, COG3523, COG3518, COG3519, COG3520 and COG3515) (82). Bacterial genomes encoding at least nine T6SS core components were regarded as harboring T6SS.

### Construction of the phylogenetic tree of bacterial genomes containing Tsi3 homologs

The DNA sequences of *rpoB* coding for DNA-directed RNA polymerase subunit beta were retrieved for bacterial genomes containing Tsi3 homologs. Sequences were clustered using CD-HIT (83) with a sequence identity threshold of 0.99. Representative sequences from the identified clusters were aligned using MAFFT v7 FFT-NS-i (84, 85). The evolutionary history was inferred using the neighbor-joining method (86) with the Jukes-Cantor substitution model (JC69). The analysis involved 474 nucleotide sequences and 3,174 conserved sites. Evolutionary analyses were conducted using the MAFFT server (https://mafft.cbrc.jp/alignment/server/), and the tree was visualized using iTOL (87).

### Construction of the phylogenetic tree of the representatives of the FGE family

A list of 40 representative FGE homologs is shown in Table S1. The list was built based on the FGE homologs identified in the original paper describing the FGE family (42) as well as the seed proteins that were used to define the FGE-sulfatase Pfam family (PF03781) (https://pfam.xfam.org/family/PF03781). The protein sequences were retrieved from NCBI and trimmed according to the conserved domain. The proteins were aligned using MUSCLE (88). The evolutionary history was inferred using the Neighbor-Joining method (86). The analysis involved 40 amino acid sequences and 136 conserved positions (95% site coverage). Evolutionary analyses were conducted in MEGA7 (89) and the tree was visualized using iTOL (87).

## Acknowledgements

This study was supported by National Institutes of Health grant R01AI125363 to M.F.F. D.S. received funding from the European Research Council (ERC) under the European Union’s Horizon 2020 research and innovation program (grant agreement No. 714224), and the Israel Science Foundation (ISFl grant No. 920/17). J.L. is funded by the Washington University Chancellor’s Graduate Fellowship. The funders had no role in this study.

## References

1. Costa TRD, et al. (2015) Secretion systems in Gram-negative bacteria: Structural and mechanistic insights. Nat Rev Microbiol 13(6):343–359.

2. Fu Y, Waldor MK, Mekalanos JJ (2013) Tn-seq analysis of vibrio cholerae intestinal colonization reveals a role for T6SS-mediated antibacterial activity in the host. Cell Host Microbe 14:652–663.

3. Sana TG, et al. (2016) Salmonella Typhimurium utilizes a T6SS-mediated antibacterial weapon to establish in the host gut. PNAS 113(34):E5044–E5051.

4. Sana TG, Lugo KA, Monack DM (2017) T6SS: The bacterial “fight club” in the host gut. PLoS Pathog 13(6):e1006325.

5. Anderson MC, Vonaesch P, Saffarian A, Marteyn BS, Sansonetti PJ (2017) Shigella sonnei Encodes a Functional T6SS Used for Interbacterial Competition and Niche Occupancy Short Article Shigella sonnei Encodes a Functional T6SS Used for Interbacterial Competition and Niche Occupancy. Cell Host Microbe 21:769–776.

6. Borgeaud S, Metzger LC, Scrignari T, Blokesch M (2015) The type VI secretion system of Vibrio cholerae fosters horizontal gene transfer. Science (80-) 347(6217):63–67.

7. Cooper RM, Tsimring L, Hasty J (2017) Inter-species population dynamics enhance microbial horizontal gene transfer and spread of antibiotic resistance. Elife 6:e25950.

8. Trunk K, et al. (2018) The type VI secretion system deploys antifungal effectors against microbial competitors. Nat Microbiol 3:920–931.

9. Hachani A, Wood TE, Filloux A (2016) Type VI secretion and anti-host effectors. Curr Opin Microbiol 29:81–93.

10. Jana B, Salomon D (2019) Type VI secretion system: a modular toolkit for bacterial dominance. Future Microbiol 14(16):1451–1463.

11. Hernandez RE, Gallegos-Monterrosa R, Coulthurst SJ (2020) Type VI secretion system effector proteins: Effective weapons for bacterial competitiveness. Cell Microbiol (22:e13241). doi:10.1111/cmi.13241.

12. Durand E, Cambillau C, Cascales E, Journet L (2014) VgrG, Tae, Tle, and beyond: The versatile arsenal of Type VI secretion effectors. Trends Microbiol 22(9):498–507.

13. Russell AB, Peterson SB, Mougous JD (2014) Type VI secretion system effectors: poisons with a purpose. Nat Rev Microbiol 12:137–148.

14. Miyata ST, Bachmann V, Pukatzki S (2013) Type VI secretion system regulation as a consequence of evolutionary pressure. J Med Microbiol 62:663–676.

15. Ting S, et al. (2018) Bifunctional immunity proteins protect bacteria against FtsZ-targeting ADP-ribosylating toxins. Cell 175:1380–1392.

16. Ahmad S, et al. (2019) An interbacterial toxin inhibits target cell growth by synthesizing (p)ppApp. Nature 575(7784):674–678.

17. Tang JY, Bullen NP, Ahmad S, Whitney JC (2018) Diverse NADase effector families mediate interbacterial antagonism via the type VI secretion system. J Biol Chem 293(5):1504–1514.

18. Whitney JC, et al. (2015) An interbacterial NAD(P)+ glycohydrolase toxin requires elongation factor Tu for delivery to target cells. Cell 163:607–619.

19. Lopez J, Feldman MF (2018) Expanding the molecular weaponry of bacterial species. J Biol Chem 293(5):1515–1516.

20. Mok BY, et al. (2020) A bacterial cytidine deaminase toxin enables CRISPR-free mitochondrial base editing. Nature 583(7817):631–637.

21. Sibinelli-Sousa S, et al. (2020) A family of T6SS antibacterial effectors related to L,D-transpeptidases targets the peptidoglycan. Cell Rep 31:107813.

22. Le N-H, Pinedo V, Lopez J, Cava F, Feldman MF (2021) Killing of Gram-negative and Gram-positive bacteria by a bifunctional cell wall-targeting T6SS effector. BioRxiv:https://doi.org/10.1101/2021.03.04.433973.

23. Robitaille S, Trus E, Ross BD (2020) Bacterial Defense against the Type VI Secretion System. Trends Microbiol DOI: 10.10. doi:10.1016/j.tim.2020.09.001.

24. Hersch SJ, Manera K, Dong TG (2020) Defending against the type six secretion system: beyond immunity genes. Cell Rep 33(2):108259.

25. Le N, et al. (2020) Peptidoglycan editing provides immunity to Acinetobacter baumannii during bacterial warfare. Sci Adv 6:eabb5614.

26. Toska J, Ho BT, Mekalanos JJ (2018) Exopolysaccharide protects Vibrio cholerae from exogenous attacks by the type 6 secretion system. PNAS 115(31):7997–8002.

27. Espaillat A, et al. (2016) Chemometric analysis of bacterial peptidoglycan reveals atypical modifications that empower the cell wall against predatory enzymes and fly innate immunity. J Am Chem Soc 138:9193–9204.

28. Hersch SJ, et al. (2020) Envelope stress responses defend against type six secretion system attacks independently of immunity proteins. Nat Microbiol 5(5):706–714.

29. Kamal F, et al. (2020) Differential cellular response to translocated toxic effectors and physical penetration by the type VI secretion system. Cell Rep 31:107766.

30. Lories B, et al. (2020) Biofilm bacteria use stress responses to detect and respond to competitors. Curr Biol 30:1–14.

31. Russell AB, et al. (2012) A widespread bacterial type VI secretion effector superfamily identified using a heuristic approach. Cell Host Microbe 11:538–549.

32. Benz J, Sendlmeier C, Barends TRM, Meinhart A (2012) Structural insights into the effector-immunity system Tse1/Tsi1 from Pseudomonas aeruginosa. PLoS One 7(7):e40453.

33. Ding J, Wang W, Feng H, Zhang Y (2012) Structural Insights into the Pseudomonas aeruginosa Type VI Virulence Effector Tse1 Bacteriolysis and Self-protection. 287(32):26911–26920.

34. Ross BD, et al. (2019) Human gut bacteria contain acquired interbacterial defence systems. Nature 575(7781):224–228.

35. Kirchberger PC, Unterweger D, Provenzano D, Pukatzki S (2017) Sequential displacement of type VI secretion system effector genes leads to evolution of diverse immunity gene arrays in Vibrio cholerae. Sci Rep 7(45133):1–12.

36. Weber BS, et al. (2016) Genetic dissection of the type VI secretion system in Acinetobacter and identification of a novel peptidoglycan hydrolase, TagX, required for its biogenesis. MBio 7(5):e01253–16.

37. Lazzaro M, Feldman MF, Vescovi EG (2017) A transcriptional regulatory mechanism finely tunes the firing of type VI secretion system in response to bacterial enemies. MBio 8:e00559–17.

38. Dar Y, Salomon D, Bosis E (2018) The Antibacterial and Anti-Eukaryotic Type VI Secretion System MIX-Effector Repertoire in Vibrionaceae. Mar Drugs 16:433.

39. Jana B, Fridman CM, Bosis E, Salomon D (2019) A modular effector with a DNase domain and a marker for T6SS substrates. Nat Commun 10(3595).

40. Fridman CM, Keppel K, Gerlic M, Bosis E, Salomon D (2020) A comparative genomics methodology reveals a widespread family of membrane-disrupting T6SS effectors. Nat Commun 11(1085). doi:10.1038/s41467-020-14951-4.

41. Ringel PD, Hu D, Basler M (2017) The Role of Type VI Secretion System Effectors in Target Cell Lysis and Subsequent Horizontal Gene Transfer. Cell Rep 21:3927–3940.

42. Landgrebe J, Dierks T, Schmidt B, Figura K Von (2003) The human SUMF1 gene, required for posttranslational sulfatase modification, defines a new gene family which is conserved from pro-to eukaryotes. Gene 316:47–56.

43. Ringel MT, Dra G, Bru T (2018) PvdO is required for the oxidation of dihydropyoverdine as the last step of fluorophore formation in Pseudomonas fluorescens. J Biol Chem 293(7):2330–2341.

44. Yuan Z, et al. (2017) Crystal structure of PvdO from Pseudomonas aeruginosa. Biochem Biophys Res Commun 484:195–201.

45. Appel MJ, Bertozzi CR (2015) Formylglycine, a post-translationally generated residue with unique catalytic capabilities and biotechnology applications. ACS Chem Biol 10(1):72–84.

46. Mougous JD, Green RE, Williams SJ, Brenner SE, Bertozzi CR (2002) Sulfotransferases and sulfatases in Mycobacteria. Chem Biol 9:767–776.

47. Buono MM, Cosma MP (2010) Sulfatase activities towards the regulation of cell metabolism and signaling in mammals. Cell Mol Life Sci 67:769–780.

48. Dierks T, et al. (2003) Multiple sulfatase deficiency is caused by mutations in the gene encoding the human Ca-formylglycine generating enzyme. Cell 113:435–444.

49. Dierks T, et al. (2005) Molecular basis for multiple sulfatase deficiency and mechanism for formylglycine generation of the human formylglycine-generating enzyme. Cell 121:541–552.

50. Carlson BL, et al. (2008) Function and structure of a prokaryotic formylglycine-generating enzyme. J Biol Chem 283(29):20117–20125.

51. Knop M, Dang Q, Jeschke G, Seebeck FP (2017) Copper is a cofactor of the formylglycine-generating enzyme. ChemBioChem 18:161–165.

52. Meury M, Knop M, Seebeck FP (2017) Structural basis for copper-oxygen mediated C-H bond activation by the formylglycine-generating enzyme. Angew Chem Int Ed 56:8115– 8119.

53. Appel MJ, et al. (2019) Formylglycine-generating enzyme binds substrate directly at a mononuclear Cu(I) center to initiate O2 activation. PNAS 116(12):5370–5375.

54. Goncharenko K V, Vit A, Blankenfeldt W, Seebeck FP (2015) Structure of the sulfoxide synthase EgtB from the ergothioneine biosynthetic pathway. Angew Chem Int Ed 54:2821–2824.

55. Stampfli AR, et al. (2019) An alternative active site architecture for O2 activation in the ergothioneine biosynthetic EgtB from Chloracidobacterium thermophilum. J Am Chem Soc 141:5275−5285.

56. Hood RD, et al. (2010) A type VI secretion system of Pseudomonas aeruginosa targets a toxin to bacteria. Cell Host Microbe 7(1):25–37.

57. Russell AB, et al. (2011) Type VI secretion delivers bacteriolytic effectors to target cells. Nature 475:343–347.

58. Brooks TM, et al. (2013) Lytic activity of the Vibrio cholerae type VI secretion toxin VgrG-3 is inhibited by the antitoxin TsaB. J Biol Chem 288(11):7618–7625.

59. Unterweger D, et al. (2014) The Vibrio cholerae type VI secretion system employs diverse effector modules for intraspecific competition. Nat Commun 5(3549). doi:10.1038/ncomms4549.

60. Alteri CJ, et al. (2017) Subtle variation within conserved effector operon gene products contributes to T6SS-mediated killing and immunity. PLoS Pathog 13(11):e1006729.

61. Dörr NCD, Blokesch M (2020) Interbacterial competition and anti-predatory behaviour of environmental Vibrio cholerae strains. Environ Microbiol 22(10):4485–4504.

62. English G, et al. (2012) New secreted toxins and immunity proteins encoded within the type VI secretion system gene cluster of Serratia marcescens. Mol Microbiol 86(4):921– 936.

63. Roeser D, et al. (2006) A general binding mechanism for all human sulfatases by the formylglycine-generating enzyme. PNAS 103(1):81–86.

64. Dickmanns A, et al. (2005) Crystal structure of human pFGE, the paralog of the Ca-formylglycine-generating enzyme. J Biol Chem 280(15):15180–15187.

65. Di Venanzio G, et al. (2019) Multidrug-resistant plasmids repress chromosomally encoded T6SS to enable their dissemination. Proc Natl Acad Sci 116(4):1378–1383.

66. Tucker AT, et al. (2014) Defining gene-phenotype relationships in Acinetobacter baumannii through one-step chromosomal gene inactivation. MBio 5(4):e01313–14.

67. Hunger M, Schmucker R, Kishan V, Hillen W (1990) Analysis and nucleotide sequence of an origin of an origin of DNA replication in Acinetobacter calcoaceticus and its use for Escherichia coli shuttle plasmids. Gene 87:45–51.

68. Lopez J, Ly PM, Feldman MF (2020) The tip of the VgrG spike is essential to functional type VI secretion system assembly in Acinetobacter baumannii. MBio 11:e02761–19.

69. Scott NE, et al. (2014) Diversity Within the O-linked Protein Glycosylation Systems of Acinetobacter Species. Mol Cell Proteomics 13(9):2354–2370.

70. Studier FW (2014) Stable expression clones and auto-induction for protein production in E. coli. Structural Genomics: General Applications, Methods in Molecular Biology, pp 17–32.

71. Wu Y, Li Q, Chen X-Z (2007) Detecting protein–protein interactions by far western blotting. Nat Protoc 2(12):3278–3284.

72. Ruiz FM, et al. (2020) Structural characterization of TssL from Acinetobacter baumannii: a key component of the type VI secretion system. J Bacteriol 202(17):e00210–20.

73. Kelley LA, Mezulis S, Yates CM, Wass MN, Sternberg MJE (2015) The Phyre2 web portal for protein modeling, prediction and analysis. Nat Protoc 10(6):845–858.

74. Yang J, et al. (2015) The I-TASSER Suite: protein structure and function prediction. Nat Publ Gr 12(1):7–8.

75. Altschul SF, et al. (1997) Gapped BLAST and PSI-BLAST: a new generation of protein database search programs. Nucleic Acids Res 25(17):3389–3402.

76. Armenteros JJA, et al. (2019) SignalP 5.0 improves signal peptide predictions using deep neural networks. Nat Biotechnol 37:420–423.

77. Käll L, Krogh A, Sonnhammer ELL (2004) A combined transmembrane topology and signal peptide prediction method. J Mol Biol 338:1027–1036.

78. Jana B, Salomon D, Bosis E (2020) A novel class of polymorphic toxins in Bacteroidetes. 3(4):1–10.

79. Sievers F, et al. (2011) Fast, scalable generation of high-quality protein multiple sequence alignments using Clustal Omega. Mol Syst Biol 7(539). doi:10.1038/msb.2011.75.

80. Crooks GE, Hon G, Chandonia J, Brenner SE (2004) WebLogo: a sequence logo generator. Genome Res 14:1188–1190.

81. Marchler-Bauer A, et al. (2017) CDD/SPARCLE: functional classification of proteins via subfamily domain architectures. Nucleic Acids Res 45:D200–D203.

82. Boyer F, Fichant G, Berthod J, Vandenbrouck Y, Attree I (2009) Dissecting the bacterial type VI secretion system by a genome wide in silico analysis: What can be learned from available microbial genomic resources? BMC Genomics 10. doi:10.1186/1471-2164-10-104.

83. Li W, Godzik A (2006) Cd-hit: a fast program for clustering and comparing large sets of protein or nucleotide sequences. Bioinformatics 22(13):1658–1659.

84. Katoh K, Misawa K, Kuma K, Miyata T (2002) MAFFT: a novel method for rapid multiple sequence alignment based on fast Fourier transform. Nucleic Acids Res 30(14):3059–3066.

85. Katoh K, Rozewicki J, Yamada KD (2019) MAFFT online service: multiple sequence alignment, interactive sequence choice and visualization. Brief Bioinform 20(4):1160– 1166.

86. Saitou N, Nei M (1987) The neighbor-joining method: a new method for reconstructing phylogenetic trees. Mol Biol Evol 4(4):406–425.

87. Letunic I, Bork P (2016) Interactive tree of life (iTOL) v3: an online tool for the display and annotation of phylogenetic and other trees. Nucleic Acids Res 44:W242–W245.

88. Edgar RC (2004) MUSCLE: multiple sequence alignment with high accuracy and high throughput. Nucleic Acids Res 32(5):1792–1797.

89. Kumar S, Stecher G, Tamura K (2016) MEGA7: molecular evolutionary genetics analysis version 7.0 for bigger datasets. Mol Biol Evol 33(7):1870–1874.

